# Genomic analysis of a synthetic reversed sequence reveals default chromatin states in yeast and mammalian cells

**DOI:** 10.1101/2022.06.22.496726

**Authors:** Brendan Camellato, Ran Brosh, Matthew T. Maurano, Jef D. Boeke

## Abstract

Up to 93% of the human genome may show evidence of transcription, yet annotated transcripts account for less than 5%. It is unclear what makes up this major discrepancy, and to what extent the excess transcription has a definable biological function, or is just a pervasive byproduct of non-specific RNA polymerase binding and transcription initiation. Understanding the default state of the genome would be informative in determining whether the observed pervasive activity has a definable function. The genome of any modern organism has undergone billions of years of evolution, making it unclear whether any observed genomic activity, or lack thereof, has been selected for. We sought to address this question by introducing a completely novel 100-kb locus into the genomes of two eukaryotic organisms, *S. cerevisiae* and *M. musculus*, and characterizing its genomic activity based on chromatin accessibility and transcription. The locus was designed by reversing (but not complementing) the sequence of the human *HPRT1* locus, including ∼30-kb of both upstream and downstream regulatory regions, allowing retention of sequence features like repeat frequency and GC content but ablating coding information and transcription factor binding sites. We also compared this reversed locus with a synthetic version of the normal human *HPRT1* locus in both organismal contexts. Despite neither the synthetic *HPRT1* locus nor its reverse version coding for any promoters evolved for gene expression in yeast, we observed widespread transcriptional activity of both loci. This activity was observed both when the loci were present as episomes and when chromosomally integrated, although it did not correspond to any of the known *HPRT1* functional regulatory elements. In contrast, when integrated in the mouse genome, the synthetic *HPRT1* locus showed transcriptional activity corresponding precisely to the *HPRT1* coding sequence, while the reverse locus displayed no activity at all. Together, these results show that genomic sequences with no coding information are active in yeast, but relatively inactive in mouse, indicating a potentially major difference in “default genomic states” between these two divergent eukaryotes.

## Introduction

There is evidence that the majority of the genome may be transcribed [1], even though coding genes account for less than 2% [2], and adding noncoding RNAs (ncRNAs) brings this to only 5% [3]. There is still debate over whether this discrepancy represents truly pervasive transcription, [4-6], or might somehow be functional. The default state of the genome may be informative in addressing this issue. A genome that is repressed, or inactive, by default would generally preclude non-specific binding of RNA polymerase and thus pervasive transcription. Conversely, a genome that is open, or active, by default would present ample opportunity for transcription machinery to bind non-specifically, leading to spurious transcription initiation. The default states may also influence the frequency with which new genes are “born” from essentially random underlying sequences, an intriguing evolutionary topic to consider [7]. The truly default state of a genome is difficult to determine due to billions of years of evolutionary pressure that has acted on the present sequences. It is thus unclear whether observed genomic states are passively present by default, or actively produced by chromatin remodelers interacting with specific sequences that may have been selected for over time. A truly default chromatin state could only be determined by observing the activity of a newly-introduced large genomic sequence that has not been selected upon.

Current techniques in synthetic genomics allow the design, assembly, and delivery of very large pieces of DNA [8, 9]. Whole genetic loci, on the scale of hundreds of kilobases, can be synthesized *de novo*, with common protocols taking advantage of the highly efficient homologous recombination pathway in the yeast *Saccharomyces cerevisiae [10-12]*. While the ability to *de novo* synthesize large DNA loci, or even entire chromosomes, allows complete freedom over the sequence of the synthetic DNA, these large DNA assemblies are usually designed to recapitulate naturally occurring sequences with relatively minor sequence modifications, even when imported from different species. However, completely novel pieces of DNA can feasibly be built, and have been to different ends [8, 9, 13]. Even large pieces of DNA recapitulating mammalian loci exist as novel DNA elements in yeast during their assembly. Large DNA molecules are also assembled for functions such as data storage, thus containing sequences that are not designed for any biological function [13]. Recently, Zhou *et al*. has demonstrated that such a piece of DNA, existing as a yeast artificial chromosome for data-storage purposes, shows evidence of active transcription. However, to date there has not been any well-controlled characterization of completely novel DNA loci in mammalian genomes, or a comparison of genomic activity for the same locus in different organismal contexts.

Our workflow for synthetic regulatory genomics involves the *de novo* assembly of large DNA loci, including an intermediate step involving *S. cerevisiae*, for delivery and characterization in a desired eukaryotic context, most often mouse embryonic stem cells (mESCs) [11, 12]. This expertise positions us to 1) design and assemble a completely novel DNA locus that does not exist in nature, and 2) characterize such a novel locus in the different genomic contexts of *S. cerevisiae* and *M. musculus*. By introducing such a novel DNA locus to both yeast and mouse cells, we can investigate and compare the default genomic state in two different eukaryotic organisms.

## Results

### Design, assembly, and delivery of a large novel DNA locus, reverse HPRT1 (HPRT1R)

In order to build a completely novel large piece of DNA, we took the reverse sequence of the human *HPRT1* locus. By using the reverse sequence (importantly, not the reverse complement), we ensured that the new locus does not contain any coding information. An important advantage to our strategy is that reversing the *HPRT1* sequence completely retains sequence features such as GC-content and repeat frequency and position, that might otherwise confound analysis. This approach also provides us with a forward/coding “control” locus, the natural *HPRT1* sequence, which we have previously shown is functionally expressed in mESCs [12]. Our two systems for investigating activity of novel DNA loci are thus both *HPRT1* and *HPRT1R* in yeast in mouse cells. We can also compare the activity of forward and reverse loci in both cell types, and can compare the activity of each locus existing episomally on a YAC and chromosomally-integrated in yeast (**Fig. 1A**).

**Figure 1.**
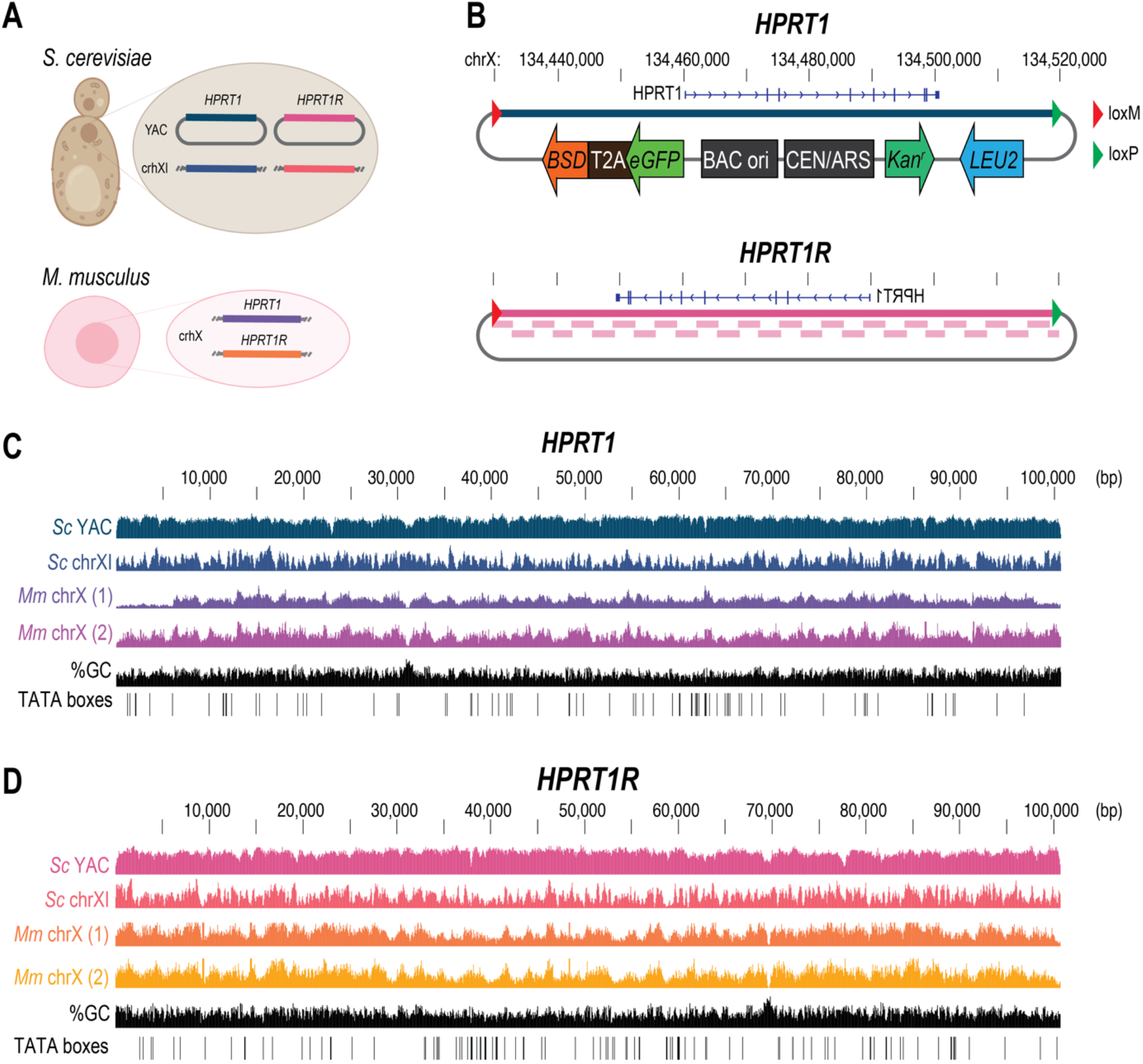
Design and construction of synthetic *HPRT1* and *HPRT1R*. **A**) Systems for interrogating *HPRT1* and *HPRT1R* activity. Episomal (YAC) and genomically integrated (chr XI) in *S. cerevisiae*, and genomically integrated (chr X) in *M. musculus*. **B**) The human *HPRT1* locus cloned into a YAC/BAC vector and flanked by lox sites for Big-In integration. *HPRT1R* was assembled *de novo* from 28 segments, shown below the locus. Vector components include *CEN/ARS* and *LEU2* for propagation and selection in *S. cerevisiae, BAC* ori and *Kanr* for propagation and selection in *E*. coli, and *eGFP-T2A-BSD* for transient selection in mammalian cells. **C, D**) NGS verification of assembled and integrated synthetic loci. SC YAC, episomal in *S. cerevisiae*; Sc chr XI, integrated in *S. cerevisiae* chr XI; Mm chr X, integrated in *M. musculus* chr X clones. (1) and (2) indicate two mESC clones. GC content (%GC) and predicted TATA boxes are shown below.

Both *HPRT1* and *HPRT1R* synthetic loci were assembled in YAC/BAC vectors allowing their Cre-mediated delivery into landing pads (LPs) pre-installed in the yeast or mouse genomes. The YAC/BAC vector contains elements for propagation and selection in both *S. cerevisiae* and *E. coli*, as well as transient selection in mammalian cells (**Fig. 1B**). The *HPRT1* locus was shuttled from a previously-assembled YAC [12] into a vector allowing Big-In delivery [11]. The synthetic *HPRT1R* locus was designed using our in-house developed software for <u>men</u>tored <u>d</u>esign <u>e</u>nvironment and LIMS (MenDEL), and assembled *de novo* into the same assembly YAC/BAC vector, first as two half-assemblies that were then combined by eSwAP-In [12] (**Fig. S1A, B**). All assemblies were verified by next-generation sequencing (**Fig. 1C, D**).

In order to integrate the synthetic loci into the yeast genome, a LP, containing a *URA3* counterselectable cassette flanked by loxM and loxP sites, was installed at *YKL162C-A*, a previously-identified safe harbor site in the yeast genome [14], in yeast strains with the *HPRT1* and *HPRT1R* assemblies as YAC/BAC episomes (**Fig. S1C**). Synthetic *HPRT1* and *HPRT1R* loci were integrated by transient expression of Cre-recombinase. Successful integrants were selected for by resistance to 5-FOA, which selects against *URA3*-expressing LP-containing yeast, verified by PCR using primers spanning the junctions between the genome and synthetic loci, and finally by next-generation sequencing.

The synthetic loci were integrated into the genome of mESCs by first installing a landing pad at the *Hprt* locus, replacing the endogenous allele on the X chromosome. This LP contains a selectable/counter-selectable marker cassette (PuroR-P2A-*PIGA*-P2A-mScarlet) flanked by loxM and loxP sites (**Fig. S1C**). LP integration was verified by PCRs spanning the genome-LP junctions and confirmed by next-generation capture-sequencing (Capture-seq) which showed both the presence of the LP as well as the absence of the native mouse *Hprt* locus. Synthetic *HPRT1* and *HPRT1R* assemblies were recovered from yeast into *E. coli* for propagation and copy number amplification, and purified for delivery to LP-containing mESCs (LP-mESCs). Purified BAC DNA was delivered to LP-mESCs by nucleofection along with a Cre-expression plasmid, and integrants were selected for the loss of the *PIGA* counterselectable marker with proaerolysin as described [11]. Integration of synthetic *HPRT1* was also selected for the presence of functional *HPRT1*, which was previously shown to be expressed in mouse cells and to complement loss of mouse *Hprt* [12]. Isolated mESC clones were screened for correct integration by junction genotyping and confirmed by next-generation Capture-seq (**Fig. 1C, D**). It was discovered through this Capture-seq verification that one mESC clone has synthetic *HPRT1* integrated in two copies (this clone is denoted as *Mm chrX (1)* throughout) (**Fig. 1C**).

### Synthetic loci are highly active in yeast

We first assessed activity of the novel synthetic loci in yeast, using three biological replicates for each of episomal or chromosomally integrated *HPRT1* and *HPRT1R*. We inspected chromatin accessibility using ATAC-seq, and observed multiple regions of highly accessible chromatin across the entire synthetic locus for both *HPRT1* and *HPRT1R* (**Fig. 2A, B**). The highly accessible regions appeared to be generally conserved across replicates and between episomal and integrated loci. Average ATAC-seq coverage depth was greater across the synthetic *HPRT1* and *HPRT1R* loci compared to the genome average in 100-kb sliding windows (**Fig. 2C**), as is visually evident when comparing the integrated synthetic loci to their surrounding native genomic regions (**Fig. S2A**). ATAC-seq coverage depth was also greater for episomal loci compared to integrated loci, even when normalizing for estimated copy number based on DNA sequencing coverage (**Fig. S1D, Fig. 2C**). The synthetic loci also contained more ATAC-seq peaks compared to the genome average (**Fig. 2D**). *HPRT1* contained 42 peaks which were shared among all three replicates, and 28 which were shared among two replicates, as an episome. Chromosomally-integrated *HPRT1* contained 12 peaks which were shared among all three replicates and 7 which were shared among two replicates. *HPRT1R* episomes contained 43 peaks which were shared among all three replicates and 39 which were shared between two replicates. Chromosomally-integrated *HPRT1R* contained 20 peaks which were shared among all three replicates and 19 which were shared among two replicates. All of these peak counts were substantially greater than the genome average of around 9 ATAC-seq peaks per 100-kb sliding window. Interestingly, both *HPRT1* and *HPRT1R* synthetic loci showed a peak of accessibility corresponding to the *HPRT1* start site or its relative position in *HPRT1R*.

**Figure 2.**
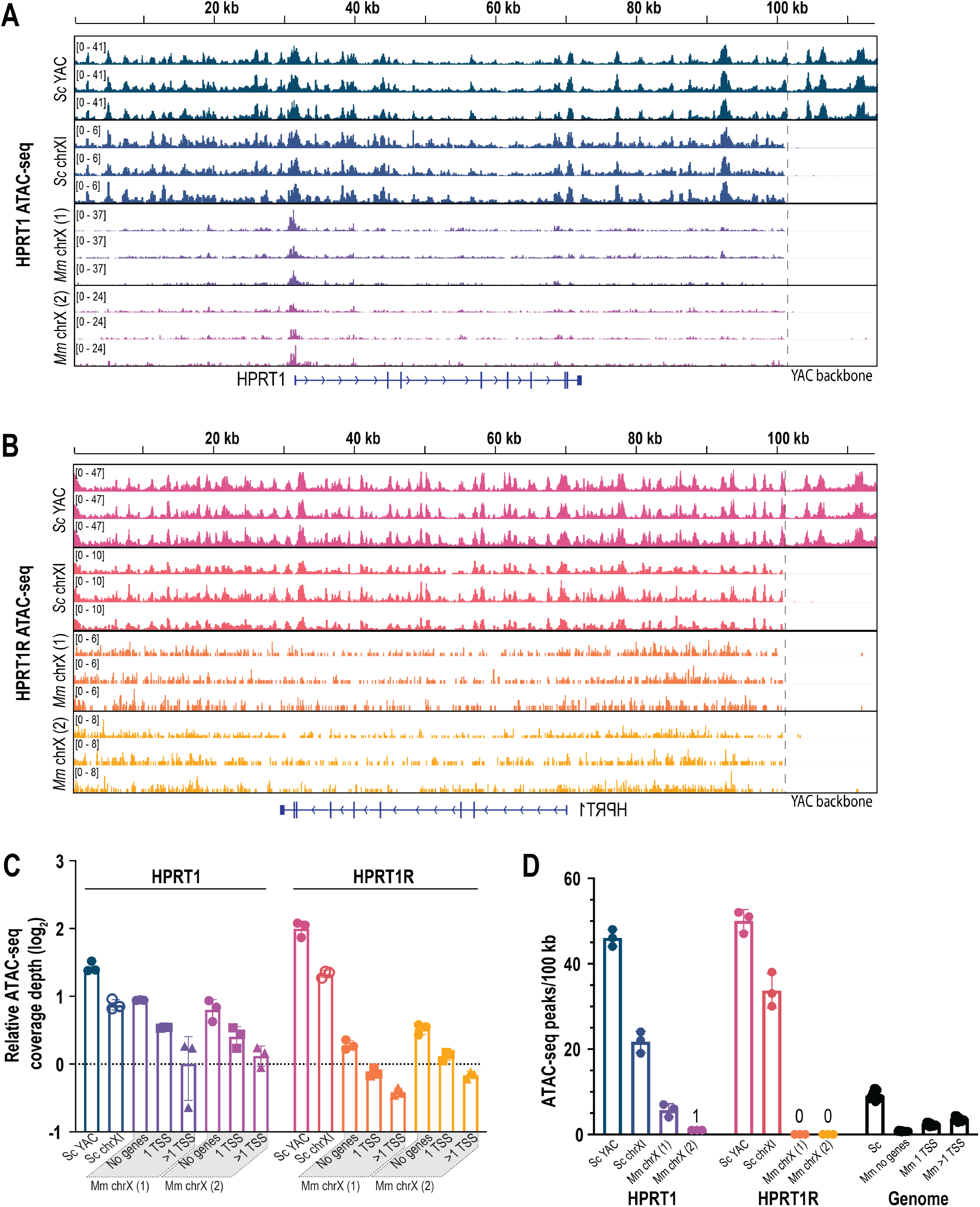
Synthetic *HPRT1* and *HPRT1R* loci are highly accessible in yeast but not in mouse ESCs. **A, B**) Browser shots of ATAC-seq reads aligned to the synthetic *HPRT1* (**A**) or *HPRT1R* (**B**) sequences. Three independent biological replicates were performed for each of *S. cerevisiae* episomal and *S. cerevisiae* chromosomal, and three replicates of each of the two clones for *M. musculus* integrations. The *HPRT1* coding sequence is shown in A, and its relative reverse position (the position in the *HPRT1R* locus corresponding to the reversed *HPRT1* coding sequence) in B. **C**) ATAC-seq read coverage depth of *HPRT1* and *HPRT1R* synthetic loci relative to the average of 100-kb sliding windows over the yeast genome (‘Sc YAC’ and ‘Sc chrXI’) or over three different regions in the mouse genome – 100-kb windows containing no genes, 1 TSS, or >1 TSS. Each biological replicate was independently compared to its own genomic averages. **D**) ATAC-seq peak counts for *HPRT1* and *HPRT1R* in each genomic context, as well as 100-kb sliding window averages for the yeast genome (Sc) or Mm10 regions with no genes, 1 TSS, or >1 TSS.

To determine whether the observed patterns of chromatin accessibility are related to transcription activity, we performed RNA-seq on episomal and chromosomally-integrated synthetic loci (**Fig. 3A, B**). Multiple RNA-seq peaks were observed for both *HPRT1* and *HPRT1R* synthetic loci. These peaks appear to be generally conserved between replicates and between episomal and chromosomal loci. Both *HPRT1* and *HPRT1R* synthetic loci showed RNA-seq coverage depth comparable to the genome average, which is gene-dense (**Fig. 3C**). Episomal *HPRT1* had 51 peaks shared by all three replicates, and 32 peaks shared by two replicates, while chromosomal *HPRT1* had 43 peaks shared by all three replicates, and 65 peaks shared by two replicates. *HPRT1R* had a total of 54 peaks shared by all three replicates, and 47 shared by two replicates, when present as an episome, and 37 peaks shared by all three replicates, and 29 shared by two replicates, when chromosomally integrated. The peaks do not appear to correspond to any known gene features in the *HPRT1* locus, and in fact they actually appear to be generally depleted around the region containing the *HPRT1* transcription unit. Interestingly, in one replicate each of episomal *HPRT1* and *HPRT1R* there was a peak corresponding to the start of the *HPRT1* gene, or its relative position in *HPRT1R*, which appears to be the only major peak that is not conserved between replicates. Relative to surrounding genomic regions, RNA expression in the *HPRT1* and *HPRT1R* loci was higher than some yeast genes, but lower than many others (**Fig. S3A**).

**Figure 3.**
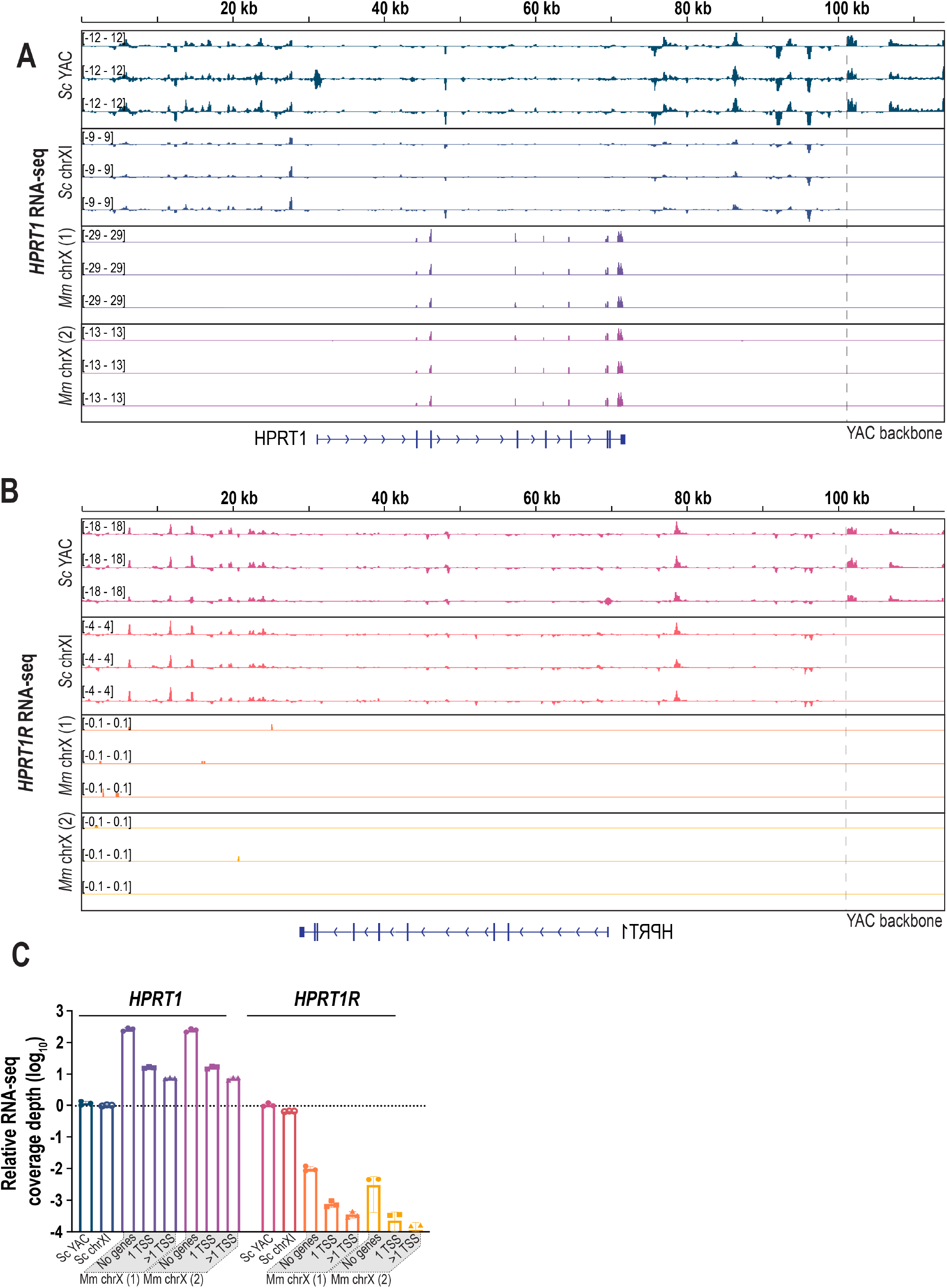
Widespread transcription of synthetic *HPRT1* and *HPRT1R* in yeast but not in mouse ESCs. **A, B**) Browser shots of RNA-seq reads aligned to the synthetic *HPRT1* (**A**) and *HPRT1R* (**B**) sequences. Three biological replicates were performed for each clone – *S. cerevisiae* episomal and genomically integrated, and two clones each for *M. musculus* integration. The *HPRT1* coding sequence is shown in **A**, and its relative reverse position (the position in the *HPRT1R* locus corresponding to the reverse *HPRT1* coding sequence) in **B. C**) RNA-seq read coverage depth of *HPRT1* and *HPRT1R* synthetic loci relative to the average of 100-kb sliding windows over the yeast genome (‘Sc YAC’ and ‘Sc chrX’) or over three different regions in the mouse genome – 100-kb windows lacking genes, or containing 1 annotated TSS, or >1 TSS. Each biological replicate was independently compared to its own genomic averages

Together, these data indicate that synthetic loci with no yeast coding information are nonetheless made accessible when present in yeast both episomally and chromosomally integrated, and peaks of accessibility generally do not correlate with known *HPRT1* functional elements. Both synthetic *HPRT1* and *HPRT1R* loci are also actively transcribed in yeast, and RNA-seq coverage depth of the synthetic loci is comparable to the gene-dense genome average.

### Synthetic loci are largely inactive in mESCs

We next performed ATAC-seq to assess the activity of the synthetic loci in mESCs. *HPRT1* showed a clear peak of accessibility at the beginning of its coding sequence (**Fig. 2A**). This agrees with data from moue early embryonic tissues [15] and a human embryonic stem line (H7-hESC) [16], both showing peaks of accessibility at the beginning of the *Hprt* and *HPRT1* coding sequences respectively (**Fig. S2C, D**). This locus showed greater coverage depth compared to 100-kb genomic windows containing no genes, one TSS, and comparable to regions containing multiple TSSs (**Fig. 2C**). Although 7, 4, and 6 ATAC-seq peaks were identified in the three replicates of clone *Mm chrX (1)*, only the peak at the *HPRT1* transcriptional start site was shared among all three replicates. The other peaks were more likely artifacts arising from the deeper coverage of this doubly-integrated clone. For clone *Mm chrX (2)*, all three replicates had only the peak at the *HPRT1* transcription start site. For the *HPRT1R* locus, there were no obvious regions of high accessibility (**Fig. 2B**). Average ATAC-seq read depth was slightly greater than genomic regions containing no genes, comparable to regions containing one TSS, and lower than regions containing multiple TSSs (**Fig. 2C**). There were no ATAC-seq peaks called for any replicates of either *HPRT1R* mESC clone.

In mESCs, only the *HPRT1* synthetic locus, unsurprisingly, showed evidence of transcription, with RNA-seq reads mapping precisely to the *HPRT1* exons. The same coverage pattern has been previously observed for the native *HPRT1* gene in a human embryonic stem cell line (H1-hESC) [17]. Coverage depth was about 2-fold greater in clone *Mm chrX (1)*, consistent with the observed 2x higher copy number. RNA-seq coverage depth was greater in the synthetic *HPRT1* locus than mouse genome regions containing no genes, one TSS, and even multiple TSSs, indicating that this human gene is highly expressed in mESCs (**Fig 3C**), but was still lower than 100-kb regions of maximal coverage depth. This elevated expression is evident when comparing depth of RNA-seq reads mapping to the *HPRT1* gene with genes in surrounding genomic regions (**Fig. S3B**). The *HPRT1R* synthetic locus demonstrated no evidence of transcription (**Fig. 3B**), and RNA-seq read depth was substantially lower than any genomic region containing one, multiple, or even no genes (**Fig. 3C**), although still greater than 100-kb regions of minimal coverage depth.

In sum, only the synthetic *HPRT1* locus was obviously active in mESCs, with a peak of accessibility corresponding specifically with the *HPRT1* start site and RNA-seq reads mapping to the *HPRT1* exons, while there was no reproducible transcriptional activity in the *HPRT1R* locus despite traces of accessibility.

### Discussion

Although the majority of the human genome may show evidence of transcription, it is still unclear whether this observed activity is functional in any way, or just a pervasive by-product of non-specific transcription initiation. As the genome has been evolving for billions of years, it is difficult to say whether any activity, or lack thereof, has been selected for over time. By introducing completely novel 100-kb DNA loci into yeast and mouse cells, we were able to assay for genomic activity in large DNA sequences that have not undergone evolutionary pressure. By comparing the activities between yeast and mouse, we were able to shed light on differences in the default genomic states between these two divergent eukaryotes. In yeast we observed widespread activity in both synthetic loci, despite not containing any promoters evolved for gene expression in yeast. Both the *HPRT1* and *HPRT1R* loci were more accessible than the rest of the yeast genome, and showed transcriptional activity comparable to the rest of this gene-dense genome. Although one of the conserved ATAC-seq peaks in yeast corresponded to the *HPRT1* transcription start site, and peaks were observed along the entire locus, the *HPRT1* coding region, and its corresponding location in *HPRT1R*, appeared more devoid of RNA-seq peaks compared to surrounding regions. It was also interesting to observe that the only clearly inconsistent peak between three replicates of episomal *HPRT1* and *HPRT1R* corresponded precisely to the *HPRT1* transcription start site, or its corresponding location in *HPRT1R*.

Recent work looking at genomic activity in a yeast episome designed for data storage similarly found that transcriptional activity could be observed throughout the entire sequence [13]. These results, along with our work here, indicate that the default genomic state in yeast is open and active chromatin, and these large synthetic loci provide ample substrates for non-specific RNA polymerase binding and transcription initiation. This may represent an evolutionary strategy in fast-growing single-celled organisms like yeast, where widespread transcription of even non-coding sequences allows a chance for new, potentially favorable genes to arise [18]. It will be interesting to follow these synthetic loci-containing strains over multiple generations and observe whether the widespread activity seen here is tuned in any way, or whether a novel gene can somehow arise from these newly-introduced sequences.

In contrast to what we observed in yeast, activity at both synthetic loci was largely restricted in mESCs. In the *HPRT1* locus, the only major peak of accessibility was at the *HPRT1* transcription start site, and RNA-seq reads only mapped to the *HPRT1* exons. This human *HPRT1* gene also appeared to be expressed at a substantially higher level than other genes in the mouse genome on average, with a greater average read depth over the 100-kb locus, although still below the level of maximally-expressed genes in this cell type. It should be noted that the current analysis did not discriminate between genes that are and aren’t expressed in mESCs, and the regions containing non-mESC genes would still be counted as *1 TSS* or *>1 TSS* regions, potentially skewing this metric. In contrast to the *HPRT1* locus, and to *HPRT1R* in yeast, the *HPRT1R* locus showed no evidence of transcriptional activity whatsoever in mESCs. There were no peaks of DNA accessibility, and any ATAC-seq coverage appeared consistent with the genome background. This suggests that the default genomic state in mouse is inactive and repressive. Although it has been argued that evidence of transcription can be found if you sequence any transcriptome deeply enough [19], here we used the rest of the genome as reference for our observations at the novel *HPRT1R* locus. The RNA-seq average read depth at this locus was substantially lower than the genomic average for 100-kb windows containing no genes, although not as low as the minimum observed depth in these regions. There are a few possible explanations for why there is detectable transcription at native geneless regions: 1) transcription associated with known and annotated genes that are not themselves located in the region of interest, either as extended transcription or from regulatory elements like enhancers [4], 2) there could be as yet unannotated functional genes, 3) it could be that low levels of transcription throughout the genome are somehow functional and thus have been selected for, for example in transcription-coupled repair or possibly other, yet undiscovered mechanism of transcription-associated genome maintenance. Pervasive transcription may also represent a historically active region in an ancestral genome, where gene expression activity has been largely but incompletely lost. We would not expect to see this kind of pervasive transcription at our synthetic *HPRT1R* locus as it has no genomic history. As with the synthetic loci in yeast, it will also be interesting to follow mESCs containing *HPRT1R* over generations to see whether any activity might arise from this locus, or through differentiation into different cell types as we have thus far only looked at undifferentiated mESCs.

In conclusion, we have used our ability to build a completely novel synthetic locus as a tool to assess the background, or default genomic activity in two eukaryotic organisms, yeast and mouse. The results indicate that the chromatin state in yeast is open and active by default, and repressed and inactive in mouse. Our results thus indicate that transcription initiation by RNA polymerase is not truly pervasive, and that any observed transcription in a large, gene-poor genome may well be functional, or maybe a remnant of previous activity in an ancestral genome.

## Materials and methods

### Design of synthetic *HPRT1R* locus

The synthetic *HPRT1R* locus was designed by taking the reverse (but not reverse-complement) sequence of the human *HPRT1* locus corresponding to hg38 chrX: 134429208-134529874. We used a software developed in-house for <u>men</u>tored <u>d</u>esign <u>e</u>nvironment and LIMS (MenDEL), to split the synthetic locus into 27 segments of ∼4-kb, and one of ∼2-kb, overlapping each other by ∼300 bp. MenDEL was also used to design primers for junction-PCR screening of the yeast harboring the assembly. Synthetic DNA segments were ordered from Qinglan Biotech, and junction PCR primers were ordered from IDT.

### Yeast assembly and BAC recovery

All yeast work was performed in the strain BY4741 using standard yeast media. For *HPRT1R* assembly, ∼50 ng each of linearized and gel-purified YAC/BAC backbone pLM1110 [11] DNA and purified assembly fragments were transformed into yeast using the high-efficiency lithium acetate method [20]. Transformants were plated on synthetic complete media lacking uracil or leucine (SC–Ura, SC–Leu) depending on the selectable marker (*URA3* for segments 1-15 half-assembly, and *LEU2* for segments 15-28 half-assembly). Successful assemblies were screened by junction PCR on crude yeast genomic DNA (gDNA) prepared from 48 colonies from each assembly. Crude yeast gDNA was prepared by performing three cycles of boiling yeast in 20 mM NaOH at 98°Cfor 3 min, followed by cooling at 4°Cfor 1 min. Junction qPCRs were set up by dispensing crude gDNA and premixed junction primer pairs into a LightCycler® 1536 Multiwell Plate (Roche 05358639001) containing 1x LightCycler® 1536 DNA Green mix (Roche 05573092001). qPCR reactions were performed using a LightCycler® 1536 Instrument (Roche 05334276001) and successful assemblies were identified based on positive results for all junctions. Candidate assemblies were verified by next-generation sequencing. Libraries were prepared from 100 ng of DNA using the NEBNext® Ultra™ II FS DNA Library Prep Kit for Illumina (NEB E7805L) with NEBNext® Multiplex Oligos for Illumina® (E7600S), according to the manufacturer’s protocol for FS DNA Library Prep Kit with Inputs ≤100 ng. Sequencing reactions were run on a NextSeq 500 system (Illumina SY-415-1001). Sequence-verified assembly YAC/BACs were recovered from yeast using the Zymoprep Yeast Miniprep I kit (Zymo Research D2001) and electroporated into TransforMax™ EPI300™ Electrocompetent *E. coli* (Lucigen EC300150), recovered in LB + 5 mM MgCl_2_ at 30°C for 1 hour and then selected on LB + kanamycin. Bacteria colonies were screened by colony PCR for one or two assembly junctions to confirm they contained the YAC/BAC, then YAC/BAC DNA was isolated from overnight cultures using ZR BAC DNA Miniprep kit (Zymo Research D4048) and verified once again by next-generation sequencing. eSwAP-In [12] was performed to combine the two *HPRT1R* half assemblies. The verified assembly YAC/BAC of segments 15-28 was purified from *E. coli* and digested with I-SceI and NotI to release the *HPRT1R* portion along with the *LEU2* marker. This digested segment was transformed into yeast harboring the YAC/BAC with segments 1-15, along with a Cas9/gRNA-expression vector with a *URA3*-targeting gRNA, pYTK-Cas9 made using the M°Clo-Yeast Toolkit [21, 22]. The Cas9-induced break in the *URA3* marker is repaired with the *HPRT1R-*15-28-*LEU2* segment using homology provided by the common segment 15 and common sequence downstream of the selection markers. eSwAP-In transformants were selected on SC–Leu and 96 colonies were picked to screen by junction PCR using a subset of primers. Candidate clones were verified by next-generation sequencing and recovered into *E. coli* as previously described.

The *HPRT1* locus was transplanted from its original assembly vector [12] by restriction digestion of purified YAC/BAC DNA with NotI and NruI to release the *HPRT1* locus, followed by co-transformation of the digested locus (∼1.5 μg) along with the new, linearized, pLM1110 assembly vector (∼100 ng) and linker DNAs that included loxP/loxM sites flanked by 200 bp of homology to the assembly vector and *HPRT1* locus (∼50 ng each). 48 colonies were picked following transformation and selection and crude yeast gDNA was screened by PCR using primers spanning the vector-*HPRT1* junctions. Candidate clones were verified by next-generation sequencing and recovered into *E. coli* as described above.

Synthetic *HPRT1* and *HPRT1R* BACs were recovered from TransforMax™ EPI300™ *E. coli* for delivery to mESCs. 250 mL cultures were grown at 30°Cwith shaking overnight in LB + kanamycin + 0.04% arabinose to induce copy number amplification of the BAC. DNA was purified using the NucleoBond Xtra BAC kit (Takara Bio 740436.25) and stored at 4°Cfor less than one week before delivering to mESCs.

### Integrating synthetic loci into the yeast genome

A landing pad (LP) containing a *URA3* cassette flanked by loxP and loxM sites was installed at YKL162C-A [14] in yeast strains harboring either *HPRT1* or *HPRT1R* episomal YAC/BACs. The LP was co-transformed along with linker DNAs with terminal homologies to the yeast genomic locus and to the LP cassette (∼200 ng each) into yeast as previously described and colonies were selected on SC–Ura. 4 colonies were picked from each transformation, screened by PCR using primers spanning the genome-LP junctions, and LP integration was verified by Sanger sequencing of PCR products spanning the genome-LP junctions. The synthetic *HPRT1* and *HPRT1R* loci were integrated by Cre-mediated recombination. A *HIS3* plasmid expressing Cre-recombinase from a galactose-inducible promoter was introduced by yeast transformation, single colonies were picked and grown to saturation in SC–His–Leu with raffinose, subcultured 1:100 in SC–His with galactose, and plated on SC+5FOA after 2 days of growth. 5FOA-resistant colonies were picked, 48 from each integration, and screened by PCR using primers spanning the yeast genome-*HPRT1/HPRT1R* junctions and finally verified by next-generation sequencing.

### mESC culture

C57BL6/6J × CAST/EiJ (BL6xCAST) Δ*Piga* mESCs, which enable PIGA-based Big-IN genome rewriting, were previously described [11]. mESCs were cultured on plates coated with 0.1% gelatin (EMD Millipore ES-006-B) in 80/20 medium comprising 80% 2i medium and 20% mESC medium. 2i medium contained a 1:1 mixture of Advanced DMEM/F12 (ThermoFisher 12634010) and Neurobasal-A (ThermoFisher 10888022) supplemented with 1% N2 Supplement (ThermoFisher 17502048), 2% B27 Supplement (ThermoFisher 17504044), 1% glutamax (ThermoFisher 35050061), 1% Pen-Strep (ThermoFisher 15140122), 0.1 mM 2-mercaptoethanol (Sigma M3148), 1,250 U/mL LIF (ESGRO ESG1107l), 3 μM CHIR99021 (R&D Systems 4423), and 1 μM PD0325901 (Sigma PZ0162). mESC medium contained knockout DMEM (ThermoFisher 10829018) supplemented with 15% FBS (BenchMark 100106), 0.1 mM 2-mercaptoethanol, 1% glutamax, 1% MEM nonessential amino acids (ThermoFisher 11140050), 1% nucleosides (EMD Millipore ES-008-D), 1% Pen-Strep, and 1,250 U/mL LIF.

### Integrating synthetic loci into the mouse genome

Integration of synthetic loci was performed using the Big-IN method [11]. First, a landing pad, LP-PIGA2, containing a polycistronic cassette - pEF1ɑ-*PuroR-P2A-PIGA-P2A-mScarlet-EF1αpA* - for selection and counterselection and flanked by loxM and loxP sites - was modified with homology arms (HAs) for targeting the LP to the mouse *Hprt* locus. Specifically, ∼130 bp HAs (amplified from a mouse *Hprt* BAC) flanked by gRNA sites for the *Hprt*-targeting gRNAs (see below) and PAMs were cloned flanking the lox sites using *Bsa*I Golden Gate Assembly. LP-PIGA2 was delivered to BL6xCAST Δ*Piga* mESCs, along with Cas9/gRNA-expression plasmids (pSpCas9(BB)-2A-GFP, Addgene #48138) expressing gRNAs that target sites flanking the *Hprt* locus to be deleted, by nucleofection using the Neon Transfection System (ThermoFisher) as described [11]. 1 million cells were used per transfection with 5 μg of the LP plasmid and 2.5 μg each of Cas9/gRNA-expression plasmids. Cells were selected with 1 μg/mL puromycin starting day 1 post-transfection, with 6-Thioguanine (6-TG, Sigma-Aldrich A4660) starting day 7 post-transfection to select for the loss of *Hprt* and with 1 μM ganciclovir (GCV, Sigma PHR1593) to select against the LP plasmid backbone that contained a HSV1-ΔTK expression cassette. Candidate clones were picked on day 10, screened by PCR using primers spanning the mouse genome-LP junctions and with primers for validating the loss of the endogenous *Hprt* gene and the absence of LP backbone or pSpCas9 plasmid integration. mESC clones were further verified by next-generation baited capture sequencing (Capture-seq, below) [11] that the *Hprt* locus was deleted and the landing pad was present on target.

Delivery of the synthetic *HPRT1* and *HPRT1R* payloads were performed as described [11] using the Amaxa 2b nucleofector (program A-23). Briefly, 5 million cells were nucleofected with 5 μg pCAG-iCre and 5 μg *HPRT1* or *HPRT1R* BAC DNA. Nucleofected mESCs were treated with 10 μg/ml blasticidin for 2 days starting 1 day post-transfection to transiently select for the presence of the synthetic BACs, and then with 2 nM proaerolysin for 2 days starting day 7 post-transfection to select for loss of *PIGA* in the LP cassette. Cells delivered with *HPRT1* were also selected with HAT medium (Thermo Fisher Scientific 21060017) starting day 7 post-transfection. Clones were picked on day 9 post-transfection, expanded, and screened first by quantitative PCR aided by an Echo 550 liquid handler (LabCyte) as described (Brosh bioRxiv 2022) using primers spanning the junctions between the mouse genome and *HPRT1* or *HPRT1R* synthetic loci, and verified by Capture-seq (see below) [11].

### Capture-sequencing

Targeted capture-sequencing (Capture-seq) was performed as previously described [11]. Biotinylated bait DNA was generated by nick translation from purified BACs and plasmids of interest: the mouse *Hprt*-containing BAC (RP23-412J16, BACPAC Resources Center), the synthetic *HPRT1* and *HPRT1R* BACs, LP-PIGA2, pCAG-iCre, and pSpCas9(BB)-2A-GFP.

### ATAC-seq sample preparation

For yeast, 3 independent clones for each strain were picked and inoculated into 5 mL of SC–Leu (for YAC/BAC strains) or YPD (for integration strains) for overnight culture at 30°C. Saturated overnight cultures were diluted 1:50 and cultured for 5 hours at 30°C, until OD_600_ reached 0.5-0.6. ∼5×10^6^ cells were taken from each culture, pelleted at 3,000 rcf for 5 minutes, washed twice with 500 μL spheroplasting buffer (1.4 M Sorbitol, 40 mM HEPES-KOH pH 7.5, 0.5 mM MgCl2), resuspended in 100 μL spheroplasting buffer with 0.2 U/μL zymolyase (Zymo Reasearch E1004), then incubated for 30 minutes at 30°Con rotator. Spheroplasts were washed twice with 500 μL spheroplasting buffer then resuspended in 50 μL 1x TD buffer with TDE (Illumina 20034197). Tagmentation was performed for 30 minutes at 37°Con a rotator and DNA was purified using the DNA Clean and Concentrator 5 kit (Zymo Research D4004). PCR was performed as previously described [23] using 11 total cycles. The libraries were sequenced with 36 bp paired-end reads on a NextSeq 500 for ∼1M reads per sample.

For mESCs, 3 independent cultures of each cell line were grown to medium confluency in 6-well plates. Cells were harvested by washing once with PBS, dissociating into single-cell suspending with TrypLE Express (ThermoFisher 12604013) and then neutralizing with mESC media. Cells were counted and 50,000 were taken for tagmentation. Cells were pelleted at 500 rcf for 5 minutes at 4°C, washed with 50 μL cold PBS, resuspended in 50 μL cold ATAC lysis buffer (10 mM Tris-HCl, pH 7.4, 10 mM NaCl, 3 mM MgCl2, 0.1% IGEPAL CA-630), spun down at 500 rcf for 10 mins at 4°C, resuspended in 50 μL TDE mix, and incubated at 37°Con rotator for 30 mins. DNA was purified using the DNA Clean and Concentrator 5 kit. PCR was performed as previously described [23] using 10 total cycles. The libraries were sequenced with 36 bp paired-end reads on a NextSeq 500 for ∼50M reads per sample.

### RNA-seq sample preparation

For yeast, the remaining culture that was not used for ATAC-seq was centrifuged at 3,000 rcf for 5 minutes to pellet cells, washed once with water, pelleted again at 3,000 rcf for 5 minutes, and then pellets were frozen at -80°C. Frozen pellets were resuspended in 200 μL lysis buffer (50 mM Tris-HCl pH 8, 100 mM NaC) and lysed by disruption with an equal volume of acid washed glass beads, vortexing 10 × 15 seconds. 300 μL lysis buffer was added and samples were mixed by inversion followed by a short centrifugation to collect all liquid in the tube. 450 μL of supernatant was mixed with an equal volume of phenol:chloroform:isoamyl alcohol, vortexed for 1 minute, and centrifuged at max speed for 5 minutes. 350 μL of aqueous layer was then mixed with an equal volume of phenol:chloroform:isoamyl alcohol, vortexed for 1 minute, and centrifuged at max speed for 5 minutes. RNA was precipitated from 300 μL of the aqueous phase by adding 30 μL of 3 M NaOAc and 800 μL of cold 99.5% ethanol, briefly vortexing, and centrifuging at max speed for 10 minutes. The pellet was rinsed with 70% ethanol and dried at room temperature before dissolving in 100 μL of RNase-free DNase set (Qiagen 79254) and incubated at room temperature for 10 minutes to remove DNA. RNA was purified using the RNeasy Plus Mini kit (Qiagen 74136) and eluted in 30 μL RNase-free water. RNA-seq libraries were prepared from 1 μg total RNA using the QIAseq FastSelect -rRNA Yeast kit (Qiagen 334217) and QIAseq Stranded RNA Library kit (Qiagen 180743) according to the manufacturer’s protocol. The libraries were sequenced on a NextSeq 500 with 75 bp paired-end reads for ∼45M reads per sample.

For mESCs, the remaining cells that were not used for ATAC-seq were pelleted at 500 rcf for 5 minutes and RNA was isolated using Qiagen RNeasy Plus Mini kit, resuspending in 350 μL buffer RLT Plus + β-mercaptoethanol, with homogenization using Qiashredder columns (Qiagen 79654). RNA-seq libraries were prepared from 1 μg total RNA using QIAseq FastSelect -rRNA HMR (Qiagen 334386) and QIAseq Stranded RNA kits (Qiag) according to the manufacturer’s protocol. The libraries were sequenced on a NextSeq 500 with 75 bp paired-end reads for ∼50M reads per sample.

### Sequencing and initial data processing

Sequencing and initial data processing were performed according to Brosh et al. with modifications [11]. Illumina libraries were sequenced in paired-end mode on an Illumina NextSeq 500 operated at the Institute for Systems Genetics. All data were initially processed using a uniform mapping pipeline. Illumina sequencing adapters were trimmed with Trimmomatic v0.39 [24]. WGS and Capture-seq reads were aligned using BWA v0.7.17 [25] to a reference genome (SacCer_April2011/sacCer3 or GRCm38/mm10), including unscaffolded contigs and alternate references, as well as independently to *HPRT1* and *HPRT1R* custom references for relevant samples. PCR duplicates were marked using samblaster v0.1.24 [26]. Generation of per base coverage depth tracks and quantification was performed using BEDOPS v2.4.35 [27]. Data were visualized using the University of California, Santa Cruz Genome Browser. The sequencing processing pipeline is available at https://github.com/mauranolab/mapping.

### DNA copy number estimation sequencing analysis

For copy number estimation in yeast strains, coverage depth was calculated for the synthetic *HPRT1* and *HPRT1R* loci as well as the entire yeast genome (excluding chrM) using samtools depth [28], and the calculated depth of the synthetic loci was divided by the genome average.

### ATAC-seq analysis

Initial sequencing data processing was performed as described above. Reads were also mapped to custom reference sequences in which the synthetic *HPRT1* and *HPRT1R* sequences were inserted at their specific integration sites in the mm10 and sacCer3 genomes, made using the *reform* tool (https://gencore.bio.nyu.edu/reform/). ATAC-seq reads were aligned using bowtie2 [29] to reference genomes (SacCer_April2011/sacCer3 or GRCm38/mm10) as well as custom references. Aligned bam files were converted to bigwig using bamCoverage (deepTools 3.5.0) [30] with bin size 1, normalized using RPGC to an effective genome size of 12000000 for sacCer3 and 2652783500 for mm10, and visualized using IGV [31]. Peaks were called using macs2 [32] with the parameters: --nomodel -f BAMPE --keep-dup all -g 1.2e7 (sacCer3)/1.87e9 (mm10). For relative coverage analysis, average coverage depth was calculated over the synthetic *HPRT1* and *HPRT1R* loci, 100-kb sliding windows of yeast genome, and 100-kb sliding windows of the mouse genome (as described below) using samtools bedcov [28], which reports the total read base count (the sum of per base read depths) per specified region, and then dividing the total read base count by the region size - 100,735 bp *HPRT1*/*HPRT1R* loci or 100-kb windows. For the yeast genome, the average of the 100-kb windows was then calculated. For the mouse genome, the average of the 100-kb windows for each category was then calculated. The average coverage depth over the synthetic loci was then divided by the relevant genome average to determine relative coverage depth in each context (i.e. *HPRT1* average coverage / average coverage of yeast 100-kb windows = relative coverage of *HPRT1* compared to the yeast genome average). For peak analysis, total peaks were counted across the *HPRT1*/*HPRT1R* loci, averaged over the yeast genome 100-kb windows, and averaged over the mouse genome 100-kb windows in each category.

For mouse genome comparative analysis, the genome was split into 100-kb sliding windows with 10-kb step size using bedtools makewindows [33]. The 100-kb sliding windows were then filtered to exclude encode blacklist regions [34] and assigned to three categories: *no genes* for windows that overlapped no annotated transcripts (Gencode comprehensive gene annotation), *1 TSS* for windows that that overlapped only one annotated transcription start site, and *>1 TSS* for windows that overlapped more than one TSS.

### RNA-seq analysis

Initial sequencing data processing was performed as described above. Reads were also mapped to custom reference sequences in which the synthetic *HPRT1* and *HPRT1R* sequences were inserted at their specific integration sites in the mm10 and sacCer3 genomes, made using the *reform* tool (https://gencore.bio.nyu.edu/reform/). *STAR* [35] was used to align reads to custom references, without providing a gene annotation file. Aligned bam files were converted to bigwig using bamCoverage (deepTools 3.5.0) [30], filtering by strand, normalizing using TMM [36], and visualized using IGV [31]. Relative coverage analysis was performed as described above for ATAC-seq analysis.

### UCSC browser data

We obtained UCSC browser data for CpG islands [37, 38], as well as the following ENCODE data [39]. ATAC-seq data from E11.5 mouse embryonic tissue, data sets ENCSR150RMQ, ENCSR273UFV, ENCSR820ACB, ENCSR012YAB, ENCSR785NEL, ENCSR382RUC, ENCSR282YTE [15]. DNaseI data from the human H7 embryonic stem cell line (H7-hESC) (GSM736638) [16]. Long RNA-seq data from the human H1 embryonic stem cells line (H1-hESC) from Cold Spring Harbor Lab (GSM758566) [17].

## Acknowledgments

This work was supported by NIH/NHGRI grant 1RM1HG009491. Figures were created with BioRender.com.

Special thanks to Hannah J. Ashe, who was instrumental in generation of Capture-seq data.

## Supplementary Figures

**Figure S1.**
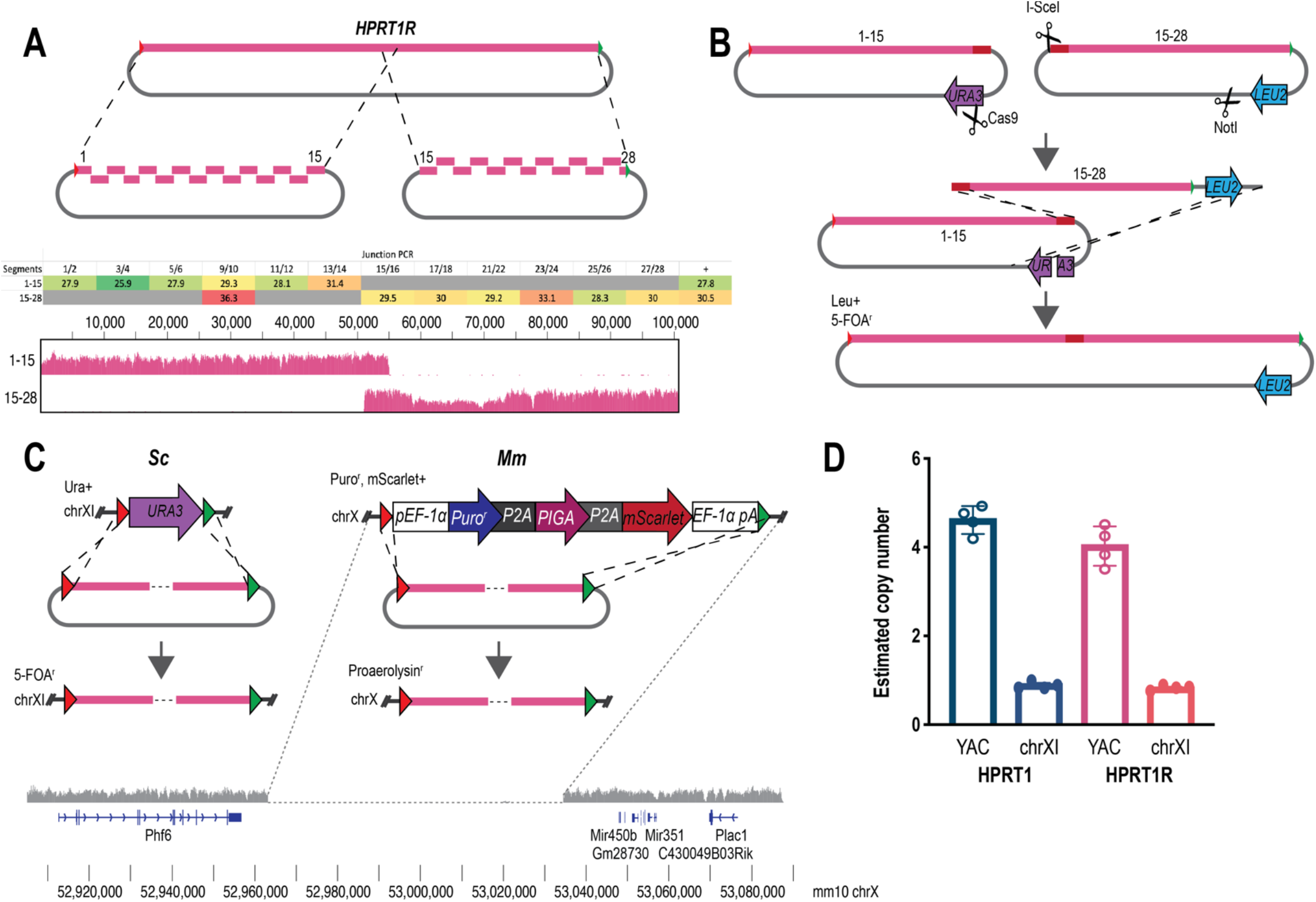
Synthetic locus assembly and cell line engineering. **A**) Assembly of *HPRT1R* in two parts, the first containing segments 1-15 and the second segments 15-28. An example junction-PCR verification is shown, reporting Ct values for qPCR reactions using primers for the indicated junctions (ie. junction 1/2 is the junction between segments 1 and 2) performed on whole yeast DNA following assembly transformation and selection. The last column represents a positive control reaction detecting a yeast genomic marker. **B**) The eSwAP-In (<u>e</u>xtrachromosomal <u>sw</u>itching <u>a</u>uxotrophies for <u>p</u>rogressive <u>in</u>tegration) strategy for combining the two *HPRT1R* half-assemblies. Segments 15-28 and the *LEU2* marker are combined with segments 1-15 and the rest of the vector backbone through homologous recombination using the common segment 15 and common sequence downstream of the selection markers as homology arms to promote recombination. Positive recombinants are selected as Leu+ and 5-FOA^r^ (Ura). **C**) Strategies for integrating synthetic loci into yeast (*Sc*) and mouse (*Mm*). For yeast, a LP containing the *URA3* selectable/counter-selectable marker was integrated at YKL162C-A on chr XI by integrative homologous recombination in strains already harboring an episomal *HPRT1* or *HPRT1R* YAC/BAC. The synthetic loci were then delivered to this locus, overwriting the *URA3* LP. For mouse, a LP containing a selectable/counterselectable cassette was installed on chrX, overwriting the endogenous *Hprt* allele. The deletion of the endogenous *Hprt* locus was verified by next-generation Capture-seq [11], as shown below. The synthetic loci were then delivered, using recombinase-mediated cassette exchange, to this locus, overwriting the LP. **D**) Estimated copy number of *HPRT1* and *HPRT1R* in yeast when existing episomally (YAC) or integrated (chrXI). Copy number was estimated based on whole genome sequencing coverage depth of the synthetic loci relative to the genome average copy number of *HPRT1* and *HPRT1R* in yeast when existing episomally (YAC) or integrated (chrXI). Copy number was estimated based on whole genome sequencing coverage depth of the synthetic loci relative to the genome average.

**Figure S2.**
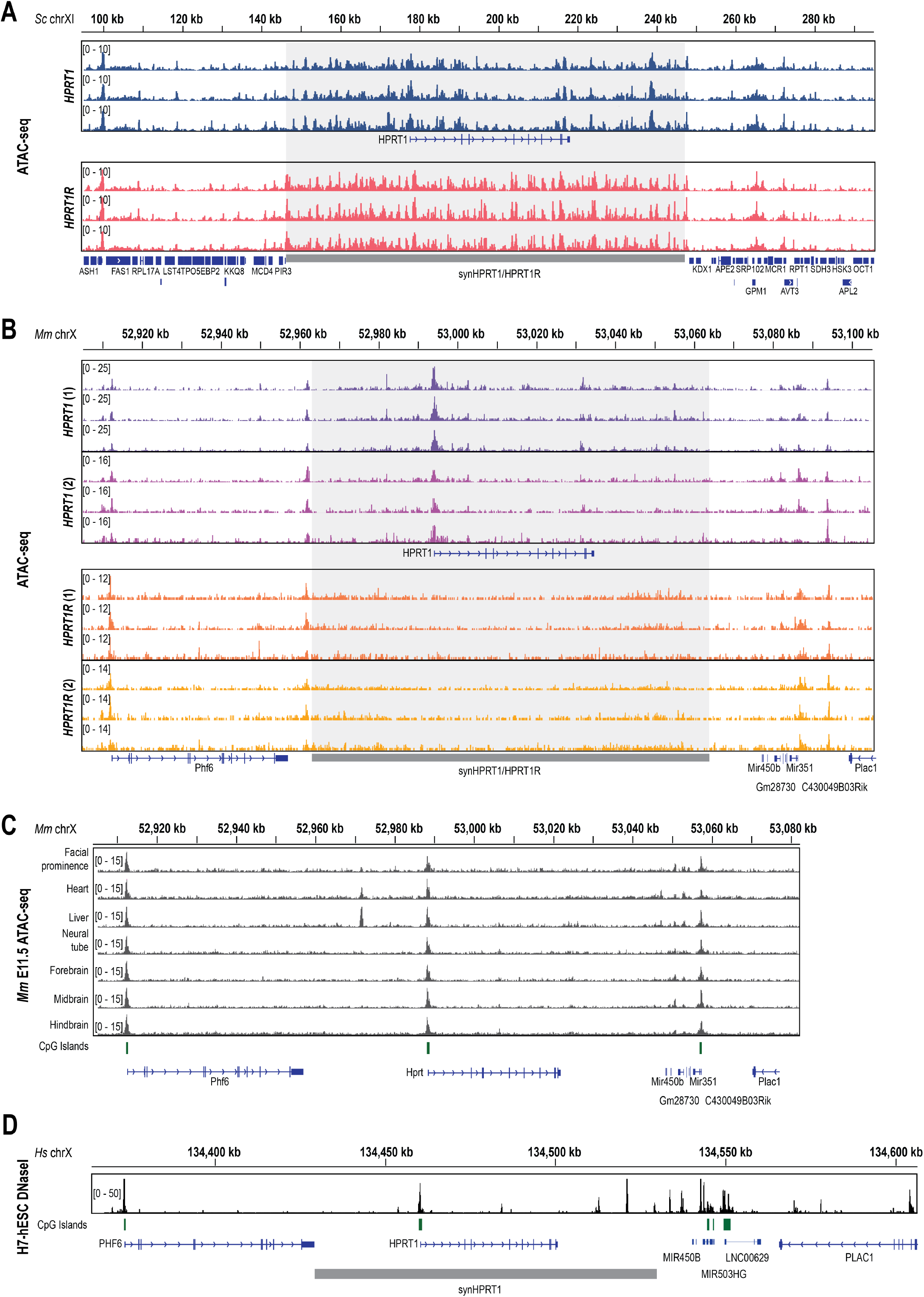
ATAC-seq of native genomic contexts. **A, B**) ATAC-seq reads were mapped to custom references in which the synthetic *HPRT1* or *HPRT1R* sequences were inserted at the corresponding positions in the yeast (**A**) or mouse (**B**) genomes. The synthetic loci are shaded, and the position of the *HPRT1* coding sequence is indicated. Endogenous genes flanking the integration sites are also shown. **C**) ENCODE ATAC-seq data in seven mouse embryonic tissues from E11.5 embryos showing coverage at the native mouse locus, including the mouse *Hprt* gene. Predicted CpG islands are shown below. **D**) ENCODE Dnase I hypersensitivity signal data showing coverage at the native human *HPRT1* locus in the H7 human embryonic stem cell line (H7-hESC). Predicted CpG islands are shown below. The region that was cloned as the synthetic *HPRT1* locus is indicated.

**Figure S3.**
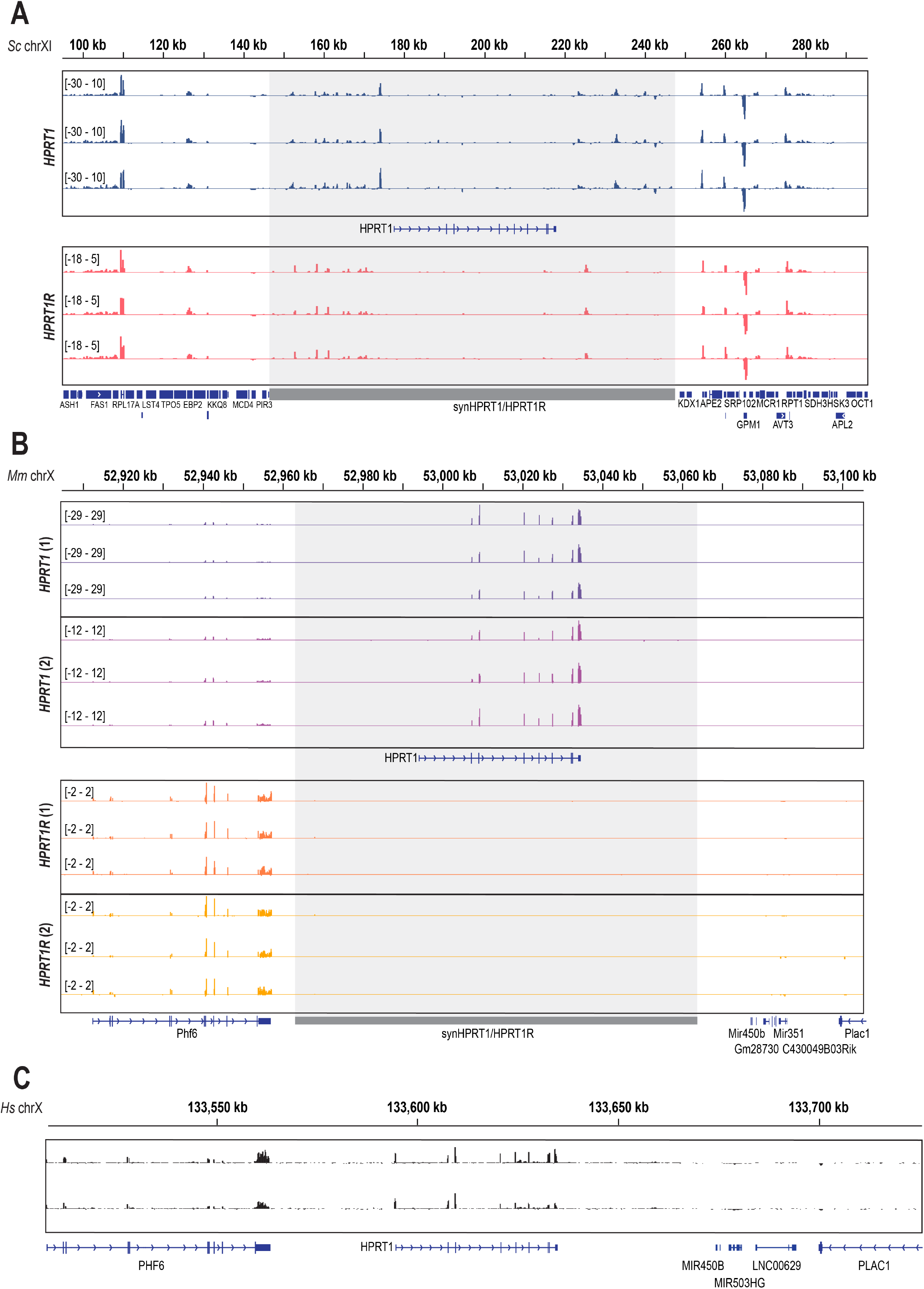
RNA-seq of native genomic contexts. **A, B**) RNA-seq reads were mapped to custom references in which the synthetic *HPRT1* or *HPRT1R* sequences were inserted at the corresponding positions in the yeast (**A**) or mouse (**B**) genomes. The synthetic loci are shaded, and the position of the *HPRT1* coding sequence is indicated. Endogenous genes flanking the integration sites are also shown. **C**) ENCODE long RNA-seq data from the H1 human embryonic stem cell line (H1-hESC) mapped to the native human *HPRT1* locus.

